# Two *Spag6* genes control sperm formation and male fertility in mice

**DOI:** 10.1101/2025.07.18.665465

**Authors:** Yunhao Liu, Wei Li, Tao Li, Cheng Zheng, Changmin Niu, Alain Schmitt, Yi Tian Yap, Mohammad Abdulghani, Shuiqiao Yuan, Christian Melander, Jerome F Strauss, Aminata Toure, Ling Zhang, Zhibing Zhang

## Abstract

Sperm-associated antigen 6 (SPAG6) is the mammalian orthologue of *Chlamydomonas* PF16, a central axonemal protein essential for flagellar motility. In mice, two homologous genes have been identified: the ancestral *Spag6* on chromosome 2 and the evolutionary derived *Spag6l* on chromosome 16. Although *Spag6* knockout mice (*Spag6^−/−^*) are phenotypically normal, the surviving *Spag6l^−/−^* males are infertile. To further investigate the roles of SPAG6 and SPAG6L, we generated compound mutants by crossing the two knockout lines. Compound heterozygous *Spag6^+/−^; Spag6l^+/−^* mice are fertile, while all *Spag6^−/−^; Spag6l^+/−^* males are infertile despite grossly normal appearance. Histological and ultrastructural analyses revealed defective spermiogenesis, including abnormal chromatin condensation, malformed acrosome and manchette, and disorganized mitochondrial and fibrous sheath. Both SPAG6 and SPAG6L bind to SPINK2, a key regulator of acrosome function, but SPAG6 has an approximately 10-fold higher binding affinity than SPAG6L. Moreover, SPAG6 modulates testicular AKAP4 and SPAG16L levels, which are critical components of the fibrous sheath and central apparatus respectively. Notably, SPAG6 suppresses tubulin acetylation, whereas SPAG6L enhances this post-translational modification, suggesting antagonistic roles in microtubule assembly. Overall, our findings demonstrate that SPAG6 and SPAG6L coordinately regulate sperm formation and male fertility during evolution.

## Introduction

Mammalian sperm-associated antigen 6 (SPAG6) is an orthologue of *Chlamydomonas reinhardtii PF16*, a protein localized within the central apparatus of the axoneme and essential for flagellar motility (1, 2). It was first identified through the screening of a human testis cDNA library using serum from an infertile man (3). Whole-exome sequencing of Pakistani families with male infertility identified a biallelic missense substitution in the *SPAG6* gene, suggesting this gene is involved in spermatogenesis (4).

The mouse *Spag6* gene was initially cloned through the screening of a cDNA library from mouse mixed germ cells using a probe with a high homology to human SPAG6 (5). Subsequently, phylogenetic analysis identified the ancestral gene, *Spag6-BC061194*, which is localized on chromosome 2 and has been renamed *Spag6* (6). The originally identified *Spag6* gene, localized on chromosome 16, is thought to have originated from gene duplication of the ancestral gene during evolution and is now termed *Spag6*-like (*Spag6l*) (5,7).

Global knockout of *Spag6l* caused approximately 50% of mutant mice to die from hydrocephalus before adulthood. Among the surviving adults, females exhibited delayed time to pregnancy, while males were infertile due to disorganization of the sperm axoneme and impaired sperm motility (2). SPAG6L is located in cytoplasmic vesicles in spermatocytes as well as in the acrosome and manchette of spermatids. It also binds to numerous proteins with diverse functions. These findings suggest that SPAG6L is a multifunctional protein that not only regulates sperm motility but also plays crucial roles in spermatogenesis by interacting with multiple protein partners in distinct cellular compartments (8). Quantitative mRNA analysis and immunolocalization studies revealed that *Spag6l* is expressed in a wide range of both ciliated and non-ciliated tissues such as testis, trachea, kidney and ovary (9). This diverse tissue distribution suggests that SPAG6L is involved in various physiological processes beyond reproduction. Indeed, in *Spag6l-*deficient mice, defects in the planar cell polarity of the inner ear was shown to result in hearing loss (10). *Spag6l* knockout mice also exhibit synapse disruption and impaired humoral immunity due to the loss of centrosome polarization and actin clearance at the synaptic cleft (11). The ciliary beat frequency, rotational polarity of ciliary axoneme and basal feet polarity of brain ependymal cells and trachea epithelial cells are also significantly disrupted in *Spag6l-*deficient mice (12). Furthermore, SPAG6L regulates cell morphology, proliferation and formation of primary cilia of mouse embryonic fibroblasts (13). It also negatively regulates neuronal migration as well as neurite growth and branching (14). These findings underscore the critical role of SPAG6L in morphogenesis.

A global *Spag6* knockout mouse model (*Spag6^−/−^*) was recently generated to explore its function. Unlike the multiple physiological defects observed in the *Spag6l^−/−^* mice, the *Spag6^−/−^* mice appeared grossly normal and fertility was unaffected in both male and female mice (7). SPAG6L and SPAG6 share 93% identity in amino acid sequences, and their predicted structures reveal nearly identical folds, consisting of eight armadillo repeats responsible for protein-protein interactions. However, some amino acid differences appear to form small clusters on the protein surface of SPAG6L and SPAG6, which may serve as putative binding sites for distinct target proteins (7). Both proteins bind to TAC1, a neurotransmitter that interacts with nerve receptors and smooth muscle cells (15). Notably, SPAG6L, but not SPAG6, binds to COPS5, a key regulator of spermatogenesis (7, 8, 16). These findings suggest that while SPAG6L and SPAG6 share some common binding partners, each may also have unique interactors. In the present study, *Spag6/Spag6l* double knockout mice were generated to investigate the coordinate roles of the encoded proteins in regulation of sperm formation and male fertility.

## Materials and Methods

### Ethics statement

All animal research was approved by the Wayne State University Institutional Animal Care with the Program Advisory Committee (Protocol number: 24-02-6561) in accordance with federal and local regulations regarding the use of non-primate vertebrates in scientific research.

### Generation of double *Spag6/Spag6l* knockout mice

Global *Spag6* and *Spag6l* knockout mice were generated previously (2,7). The *Spag6l^+/−^* mice were crossed to the *Spag6^−/−^* mice, and the resulting *Spag6l^+/−^; Spag6^+/−^* males and females were crossed to each other. Genotyping was conducted using the primers described previously (2,7).

### Spermatozoa counting

Sperm cells were collected from cauda epididymides in warm PBS solution and fixed with 2% formaldehyde at room temperature for 10 min. Sperm were washed and resuspended in PBS. The number of sperm was counted using a hemocytometer chamber, and was calculated by standard methods (17).

### Spermatozoa motility assay

Sperm were collected after swimming out from the cauda epididymides in warm PBS. Sperm motility was observed using an inverted microscope (Nikon, Tokyo, Japan) equipped with 10 x objective. Movies were recorded at 15 frames/sec with a SANYO (Osaka, Japan) color charge-coupled device, high-resolution camera (VCC-3972) and Pinnacle Studio HD (version 14.0) software. For each sperm sample, 10 fields were analyzed. Individual spermatozoa were tracked using Image J (National Institutes of Health, Bethesda, MD) and the plug-in MTrackJ. Sperm motility was calculated as curvilinear velocity (VCL), which is equivalent to the curvilinear distance (DCL) traveled by each individual spermatozoon in 1 s (VCL = DCL/t).

### Fertility assessment

To test male fertility, adult mutant and control males (2-3 months old) were paired with adult wild-type females (3-4 months old) for at least two months. Mating cages typically consisted of one male and one female. The females were checked for the presence of vaginal plugs and pregnancy. Once pregnancy was detected, the females were put into separate cages. The numbers of pregnant mice and offspring from each pregnancy were recorded.

### Histology

Testes and epididymides of 3-4-month-old mice were fixed in 4% formaldehyde solution in PBS, paraffin embedded, and 5μm sections were stained with hematoxylin and eosin using standard procedures (7). Histology of these tissues was assessed using a BX51 Olympus microscope (Olympus Corp., Melville, NY; Center Valley, PA), and photographs were taken with a ProgRes C14 camera (Jenoptik Laser, Jena, Germany).

### Transmission electron microscopy

Mouse testes and epididymal spermatozoa were fixed following established protocols (18). Briefly, the samples were fixed in 0.1 M phosphate buffer (pH 7) supplemented with 3% glutaraldehyde (Grade I; Sigma-Aldrich) for 2 h at room temperature, washed in PBS and resuspended in 0.2M sodium cacodylate buffer. The samples were then post-fixed in 1% osmium tetroxide (Electron Microscopy Sciences), dehydrated and embedded in Epon resin (Polysciences Inc.). Ultra-thin sections (90 nm) were cut with a Reichert Ultracut S ultramicrotome (Reichert-Jung AG) and then stained with uranyl acetate and lead citrate prior to observation. A JEOL 1011 electron microscope (Jeol Ltd; Tokyo, Japan) was used to examine the sections and Digital Micrograph software coupled to a Gatan Erlangshen CCD camera allowed the acquisition of images.

### Immunofluorescence analysis of testis sections

Immunofluorescence analysis of mouse testes was performed as previously reported (8). Briefly, testes were fixed with 4% paraformaldehyde in 0.1M PBS (pH 7.4), and 5µm paraffin sections were made. The slides were blocked with 10% goat serum and then incubated with the indicated primary antibodies at 4°C overnight. After washing with PBS, the slides were incubated with Alexa Fluor® 488-conjugated anti-mouse IgG (Abcam, ab150113) or Alexa Fluor® 555-conjugated anti-rabbit IgG (Abcam, ab150078) secondary antibodies for 1 h at room temperature. The slides were washed with PBS and mounted using VectaMount with DAPI. Images were captured by confocal laser-scanning microscopy (Leica SD600, Leica Microsystems, Wetzlar, Germany).

### Western blot analysis

Mouse testes were lysed in radioimmunoprecipitation assay (RIPA) buffer and protein concentration was determined using the Pierce™ BCA Protein Assay Kit (Thermo Fisher Scientific, 23225). Equal amounts of protein were heated to 95°C in sample loading buffer for 5 min and resolved by SDS-PAGE. After blocking in TBS-T buffer (20 mM Tris-HCl pH 7.4, 0.15 M NaCl, 5% non-fat dry and 0.1% Tween-20) for 1 h, the membrane was incubated with indicated antibodies: SPINK2 (1:1000, GeneTex, GTX32051); Acetyl-tubulin (1:5000, Proteintech, 66200-1-Ig); SPAG16 (#3204, 1:2000, generated by our own laboratory); AKAP4 (1:4000, a gift from Dr George Gerton at University of Pennsylvania); β-actin (1:20000, Proteintech, 66009-1-Ig) at 4°C overnight. After washing with TBS-T buffer, the membrane was incubated with a horseradish peroxidase conjugated goat anti-rabbit/mouse IgG antibody at room temperature for 1 h. The proteins were detected using Pico Ultra ECL Western Blotting Detection Solution Kits (Lamda Biotech, G075).

### Direct yeast two-hybrid assay

SPAG6/pGBK-T7, SPAG6L/pGBK-T7 and SPINK2/pGAD-T7 plasmids were constructed previously (7, 8). The plasmids were transformed into AH109 yeast and direct yeast two-hybrid assay was performed as previously reported (7).

### Luciferase complementation assay

The coding sequences (CDS) of *Spag6, Spag6l* and *Spink2* were obtained by PCR and cloned into N-Luc or C-Luc vectors (supplied by Dr. James G. Granneman, Wayne State University), respectively. The primers used for cloning are listed in **Supplemental Table 1**. All the constructs (N-Luc/*Spag6*, *Spag6/*C-Luc, N-Luc/*Spag6l*, *Spag6l/*C-Luc, N-Luc/*Spink2*, *Spink2/*C-Luc) were confirmed by sequencing analysis. The resulting plasmids were co-transfected into HEK293 cells for 48 hrs. The cells were harvested, washed with PBS and resuspended in ice-cold IB buffer (10 mM HEPES, 135 mM KCl, 60 mM NaCl, 1 mM MgCl_2_). Cell lysate was extracted by sonication and distributed in 96-well plates, Luciferase activities were measured as described previously (19, 20) and readings were recorded using a Veritas microplate luminometer. Experiments were performed three times independently, and the results are presented with standard errors.

## Results

### *Spag6^−/−^; Spag6l^+/−^* males are infertile

Global *Spag6* and *Spag6l* knockout mice were generated previously (2,7). The *Spag6l^+/−^* mice were crossed to the *Spag6^−/−^* mice, and the resulting *Spag6l^+/−^; Spag6^+/−^* males and females were crossed each other to yield offspring with nine possible genotypes listed in **Supplemental Table 2**.

All the *Spag6^−/−^_;_ Spag6l^+/−^* mice survived to adulthood with no gross abnormalities (**Figure S1**). However, when the *Spag6^−/−^_;_ Spag6l^+/−^* males were crossed to the *Spag6^+/−^_;_ Spag6l^+/−^* females, none became pregnant. Given that all the *Spag6^+/−^ Spag6l^+/−^* mice were fertile during the study, we further crossed the *Spag6^−/−^_;_ Spag6l^+/−^* males to the 3-4-month-old wild-type females, and none of the females were pregnant. The *Spag6^+/+^_;_ Spag6l^+/+^* and *Spag6^+/−^_;_ Spag6l^+/−^* males were fertile, producing litters of normal size, and the testis/body weight ratio was normal (**Table 1**). Fertility of *Spag6^−/−^_;_ Spag6l^+/−^* females was also examined. All were fertile and the litter sizes were comparable with that of the *Spag6^+/−^_;_ Spag6l^+/−^* mice. However, it took a longer time to deliver the pups after the mice were caged. During the study, no *Spag6^−/−^; Spag6l^−/−^* mice survived more than three weeks, probably due to hydrocephalus, even though they appeared to be normal within one week after birth. The mice with normal fertility (five genotypes in **Supplemental Table 2**) were used as controls in this study.

**Table 1.** Fertility, fecundity and testis weight of control and *Spag6^−/−^; Spag6l^+/−^* mice.

| Genotype | Fertility<sup>a</sup> | Litter size | Testis weight (mg) | Body weight (g) | Weight ratio (Testis/Body) |
| --- | --- | --- | --- | --- | --- |
| *Spag6*<sup>+/+</sup>; *Spag6l*<sup>+/-</sup> | 6/6 | 7.8±1.2 | 200±8.60 | 19.8±0.69 | 10.09±0.39 |
| *Spag6*<sup>+/-</sup>; *Spag6l*<sup>+/-</sup> | 6/6 | 7.3±1.0 | 193±7.31 | 19.1±1.00 | 9.90±0.25 |
| *Spag6*<sup>-/-</sup>; *Spag6l*<sup>+/-</sup> | 0/6* | 0* | 182±13.4 | 19.6±1.13 | 9.30±1.31 |

<sup>a</sup> the number of fertile mice/total number of mice
\**P*<0.05

### Abnormal sperm morphology, reduced sperm concentration and motility in *Spag6^−/−^_;_ Spag6l^+/−^* mice

To determine the etiological factors contributing to male infertility in the *Spag6^−/−^_;_ Spag6l^+/−^* mice, physiological parameters of spermatozoa collected from the cauda epididymides of mice with the different genotypes were analyzed. Notably, light microscopic examination revealed that sperm density in the control (*Spag6^+/−^_;_ Spag6l^+/−^*) mice was significantly higher than those in the *Spag6^−/−^_;_ Spag6l^+/−^* mice (**Figure S2**). Sperm analysis revealed that both sperm count and proportion of morphologically normal spermatozoa were significantly lower in the *Spag6^−/−^_;_ Spag6l^+/−^* mice than in the *Spag6^+/−^_;_ Spag6l^+/−^* mice (**Figure 1a, b**). While over 70% of spermatozoa from the *Spag6^+/−^_;_ Spag6l^+/−^* mice maintained normal motility, approximately 90% of spermatozoa from the *Spag6^−/−^_;_ Spag6l^+/−^* mice were immotile (**Figure 1c, d**) and displayed multiple morphologic defects, including shortened/coiled tails, vacuoles in the midpiece, irregular flagellar thickness, and misshapen heads (**Figure 1e**).

**Figure 1.**
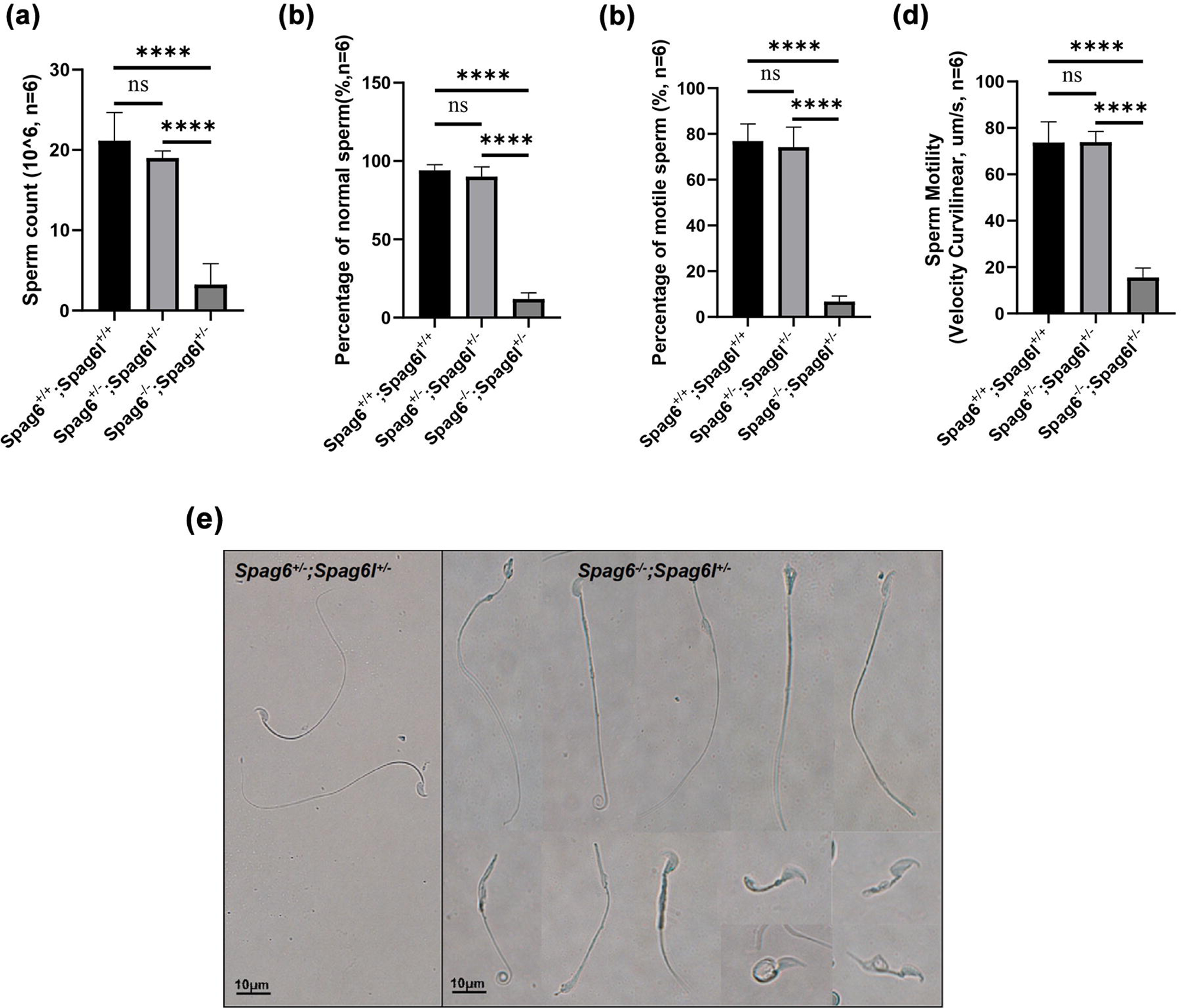
Significantly reduced sperm number, motility, and abnormally formed sperm in *Spag6^−/−^_;_ Spag6l^+/−^* mice. **(a)** The sperm number was significantly reduced in the *Spag6^−/−^_;_ Spag6l^+/−^* mice; **(b)** Significantly decreased percentage of normal sperm in the *Spag6^−/−^_;_ Spag6l^+/−^* mice; **(c, d)** Percentage of motile sperm and sperm motility was reduced in the *Spag6^−/−^_;_ Spag6l^+/−^* mice; **(e)** Morphological examination of epididymal sperm by light microscopy at high magnification. Statistically significant differences (\*\*\*\**P* < 0.001).

### Spermatogenic defects in the *Spag6^−/−^_;_ Spag6l^+/−^* mice

Testis and epididymis histology of both control (*Spag6^+/−^_;_ Spag6l^+/−^*) and *Spag6^−/−^_;_ Spag6l^+/−^* mice were examined by PAS staining. The seminiferous tubules in both groups displayed normal architecture, as reflected by the presence of all germ cell types including spermatogonia, spermatocytes, and round/elongating spermatids. However, some elongating spermatids displayed T-shaped or arrowhead-like nuclei in the stage XII-VI tubules of *Spag6^−/−^_;_ Spag6l^+/−^* mice. At stage VIII, spermatids from *Spag6^+/−^_;_ Spag6l^+/−^* mice reached maturity with heads lining the lumen, whereas some *Spag6^−/−^_;_ Spag6l^+/−^* sperm arrested in the middle layer of the seminiferous epithelium, positioned closer to the basement membrane. At stage IX, mature spermatids were released into the tubule lumen of *Spag6^+/−^_;_ Spag6l^+/−^* mice, while mislocalized spermatids were observed in the middle epithelial layer of *Spag6^−/−^_;_ Spag6l^+/−^* mice (**Figure 2a**). Furthermore, consistent with the decrease in epididymal sperm count, histological analysis showed that the epididymal lumen of *Spag6^+/−^_;_ Spag6l^+/−^* mice was filled with sperm, while significantly fewer sperm were present in the *Spag6^−/−^_;_ Spag6l^+/−^* mice (**Figure 2b**). These results suggest impaired spermiogenesis in the *Spag6^−/−^_;_ Spag6l^+/−^* mice.

**Figure 2.**
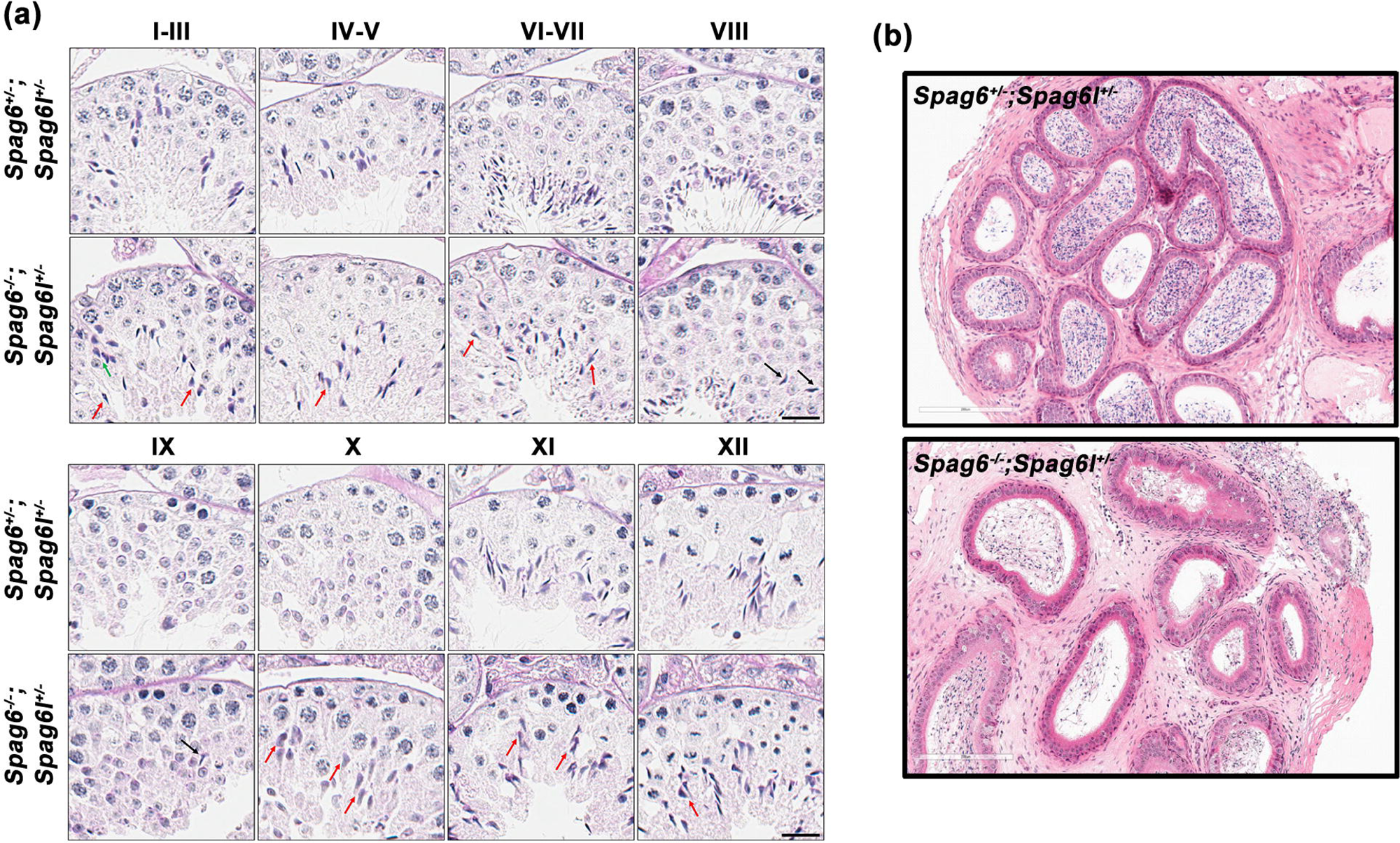
Histological examination of mouse testis and epididymis. **(a)** Periodic acid-Schiff (PAS) staining of seminiferous epithelium in mouse testis. The *Spag6^−/−^_;_ Spag6l^+/−^* mice exhibited aberrant spermatogenesis, with marked abnormalities in elongated spermatids in stages IX-X, including excessive nuclear elongation. At stage XI, “T-shaped” and “arrowhead-shaped” elongated spermatid nuclei were visible. In stages I-VII, elongated spermatids with abnormal nuclear morphology showed delayed central translocation compared to the *Spag6^+/−^_;_ Spag6l^+/−^* mice, along with misoriented nuclei. In stages VIII-IX, gradual central migration of spermatids was observed, with some retained in the middle layer of the seminiferous epithelium. Black scale bar: 20 μm. Elongated spermatids with abnormal polarity (green arrows); elongated spermatids with nuclear morphological defects (red arrows); delayed-release elongated spermatids (black arrows). **(b)** Histology of the epididymis from the *Spag6^+/−^_;_ Spag6l^+/−^* and *Spag6^−/−^_;_ Spag6l^+/−^* mice. Cauda epididymis from the *Spag6^−/−^_;_ Spag6l^+/−^* mice showing a low concentration of sperm in the lumen, compared to the *Spag6^+/−^_;_ Spag6l^+/−^* mice.

### Abnormal acrosome and manchette formation in *Spag6^−/−^_;_ Spag6l^+/−^* mice

Given the morphological defects observed in elongated spermatids from the *Spag6^−/−^_;_ Spag6l^+/−^* mice, we assessed acrosome biogenesis with fluorescently labeled PNA. No notable differences in nuclear morphology or acrosome formation were observed in step 1 to step 8 spermatids between the *Spag6^+/−^_;_ Spag6l^+/−^* and *Spag6^−/−^_;_ Spag6l^+/−^* mice. At steps 9-10, the *Spag6^−/−^_;_ Spag6l^+/−^* spermatids displayed pronounced acrosomal abnormalities, characterized by restriction of the acrosome to the apical tip of the sperm nucleus, whereas the acrosome extended laterally along the nuclei in the *Spag6^+/−^_;_ Spag6l^+/−^* spermatids. Concurrently, the nuclei of *Spag6^−/−^_;_ Spag6l^+/−^* spermatids appeared more elongated than those in the *Spag6^+/−^_;_ Spag6l^+/−^* spermatids (**Figure 3**). At steps 11~12, *Spag6^+/−^_;_ Spag6l^+/−^* spermatids showed progressive acrosomal expansion along the nuclear surface, whereas the acrosome of *Spag6^−/−^_;_ Spag6l^+/−^* spermatids maintained the apical localization. At steps 13~16, *Spag6^+/−^_;_ Spag6l^+/−^* spermatids achieved progressive nuclear compaction, with the acrosome covering the anterior and lateral portion of the sperm head. In contrast, the acrosome of *Spag6^−/−^_;_ Spag6l^+/−^* spermatids remained restricted to the apical tip, overlaying an arrowhead-shaped nucleus (**Figure 3**).

**Figure 3.**
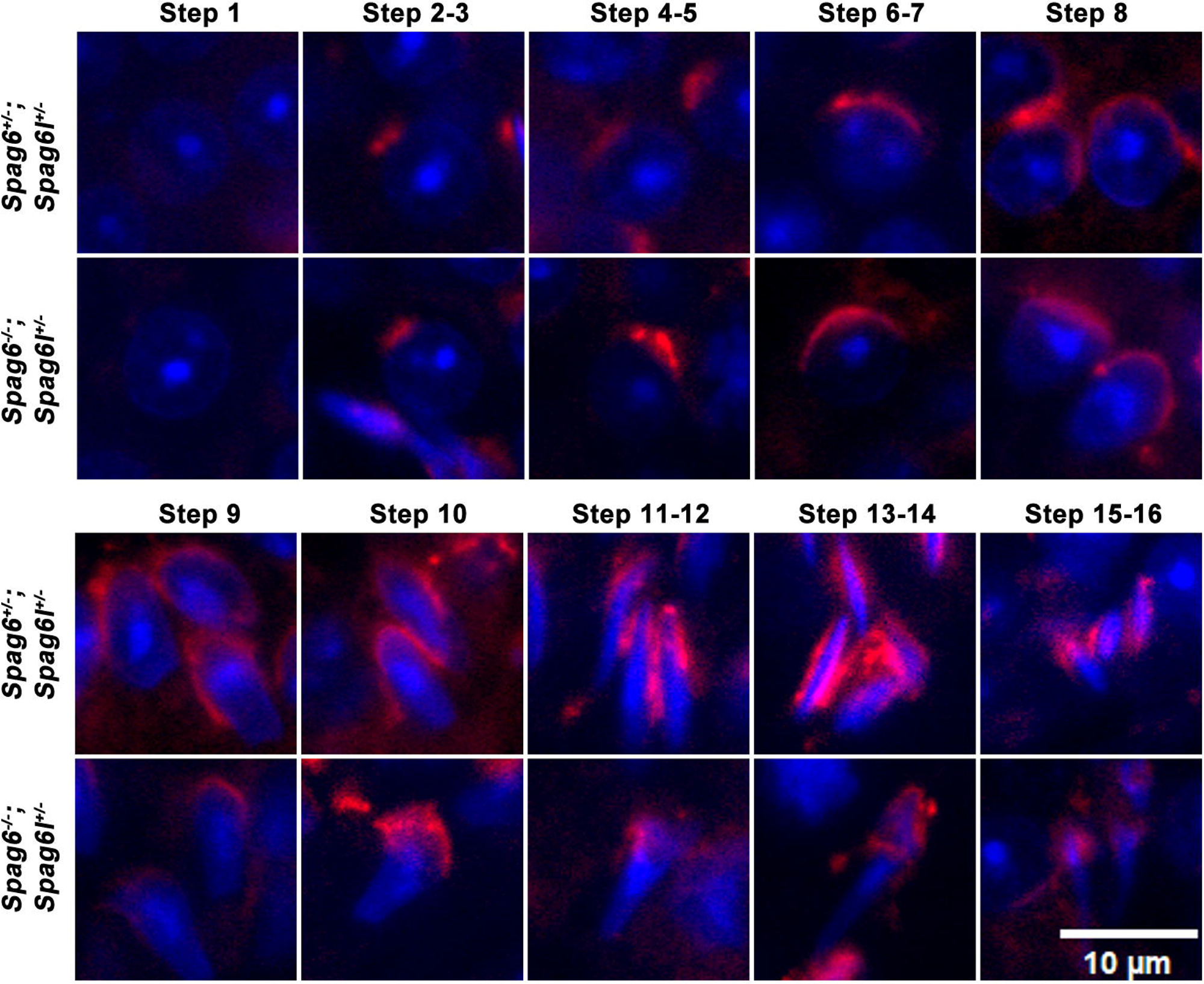
Abnormal acrosome biogenesis in the *Spag6^−/−^_;_ Spag6l^+/−^* mice. Testicular cells of *Spag6^+/−^_;_ Spag6l^+/−^* and *Spag6^−/−^_;_ Spag6l^+/−^* mice were stained with the acrosome marker peanut-lectin PNA (red). In steps 1-8 spermatids, there was no significant differences in acrosome morphology between the *Spag6^+/−^_;_ Spag6l^+/−^* and *Spag6^−/−^_;_ Spag6l^+/−^* mice. The step 9-10 spermatids from *Spag6^−/−^_;_ Spag6l^+/−^* mice displayed malformed acrosome specifically localized to the apical region of the nuclei. The step 11-12 spermatids of *Spag6^+/−^_;_ Spag6l^+/−^* mice exhibited continuous acrosomal development and progressive expansion across the nuclear surface, while *Spag6^−/−^_;_ Spag6l^+/−^* spermatids retained malformed acrosomes in the apical region of the nuclei. In steps 13-14, *Spag6^−/−^_;_ Spag6l^+/−^* spermatids have arrowhead-shaped acrosome, covering the apical tip of abnormally elongated nuclei. Nuclei were stained with DAPI (blue).

Manchette formation was assessed by immunofluorescence staining of testicular sections with α-tubulin. In both *Spag6^+/−^_;_ Spag6l^+/−^* and *Spag6^−/−^_;_ Spag6l^+/−^* mice, the manchette was first observed in step 8 spermatids at stage VIII, and disassembled in step 15 spermatids at stage IV-VI. No apparent morphological abnormalities were observed in the manchette of the *Spag6^−/−^_;_ Spag6l^+/−^* spermatids during stages VIII-IX; however, the manchette appeared more elongated and exhibited increased immunofluorescence compared to the *Spag6^+/−^_;_ Spag6l^+/−^* spermatids during stage X-XII (**Figure 4**). These findings indicate functional abnormalities in the acrosome and manchette of elongating spermatids in the *Spag6^−/−^_;_ Spag6l^+/−^* mice.

**Figure 4.**
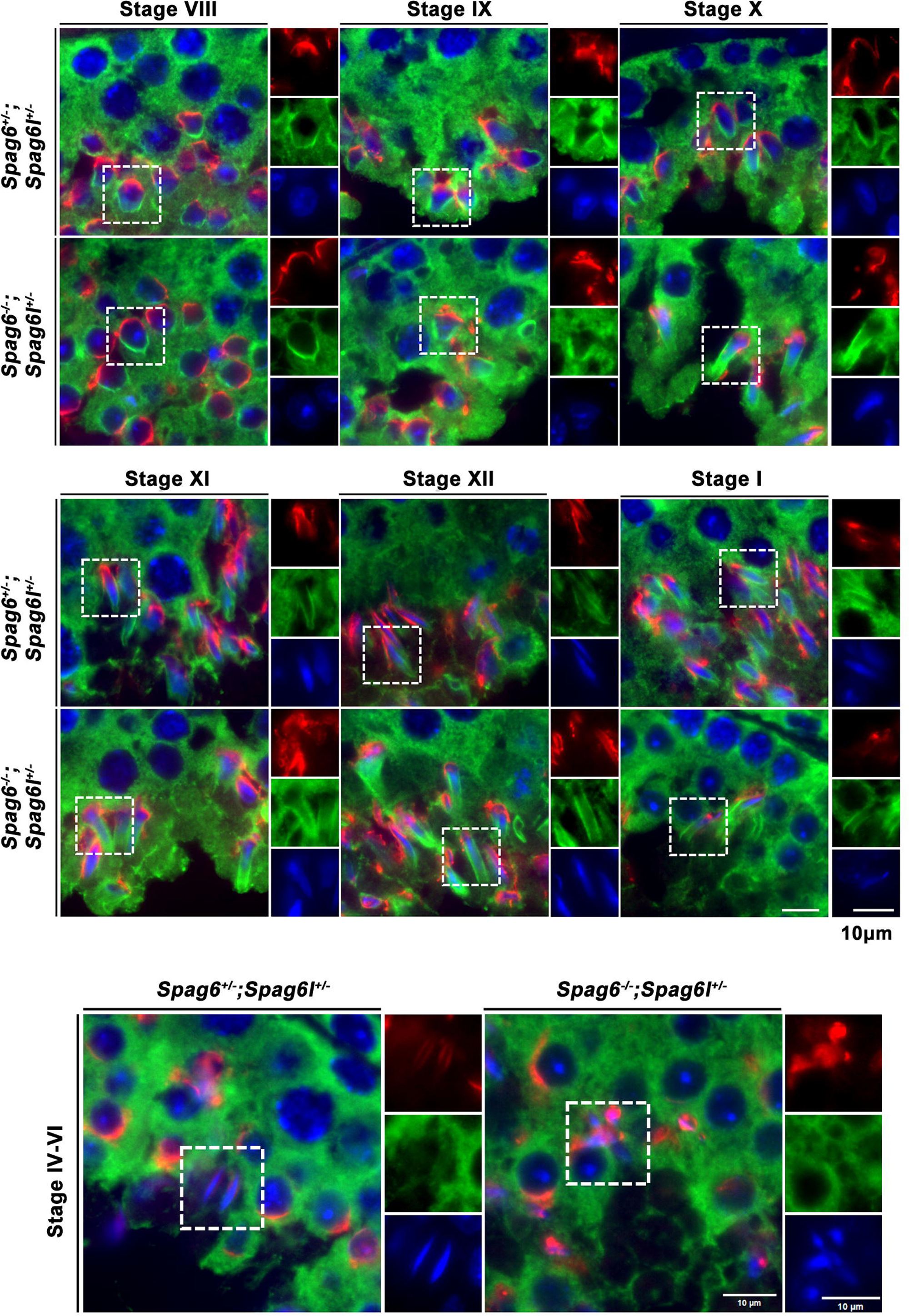
Abnormal manchette formation in the *Spag6^−/−^_;_ Spag6l^+/−^* mice. Testicular cells of *Spag6^+/−^_;_ Spag6l^+/−^* and *Spag6^−/−^_;_ Spag6l^+/−^* mice were stained with the manchette marker α-tubulin (green) and acrosome marker peanut-lectin PNA (red). In both *Spag6^+/−^_;_ Spag6l^+/−^* and *Spag6^−/−^_;_ Spag6l^+/−^*mice, the manchette was first observed in step 8 spermatids at stage VIII, and disassembled in step 15 spermatids at stage IV-VI. The manchette of *Spag6^−/−^_;_ Spag6l^+/−^* spermatids displayed no obvious morphological abnormalities in stages VIII-IX; however, it appeared more elongated in stage X-XII, compared to the *Spag6^+/−^_;_ Spag6l^+/−^* mice.

### Abnormal sperm ultrastructure in *Spag6^−/−^_;_ Spag6l^+/−^* mice

To examine ultrastructural changes in epididymal sperm, TEM analysis was performed on the axoneme and peri-axonemal structures of sperm flagella. In *Spag6^+/−^_;_ Spag6l^+/−^* mice, the sperm heads exhibited normal morphology with intact acrosomes covering the nuclear surface. The sperm flagella displayed the canonical “9+2” axoneme and normally organized outer dense fibers (ODF), mitochondrial and fibrous sheaths. In contrast, multiple abnormalities were discovered in spermatozoa collected from the *Spag6^−/−^_;_ Spag6l^+/−^* mice. The sperm head contained an abnormally elongated nucleus, malformed acrosome and excess cytoplasmic remnants. The flagellar midpiece displayed partial or complete loss of the “9+2” microtubule arrangement and disorganized outer dense fibers. Additionally, the principal piece showed a distorted fibrous sheath. Similar axonemal abnormalities were observed in the principal and end pieces of the sperm flagella (**Figure 5a, Figure S3**).

**Figure 5.**
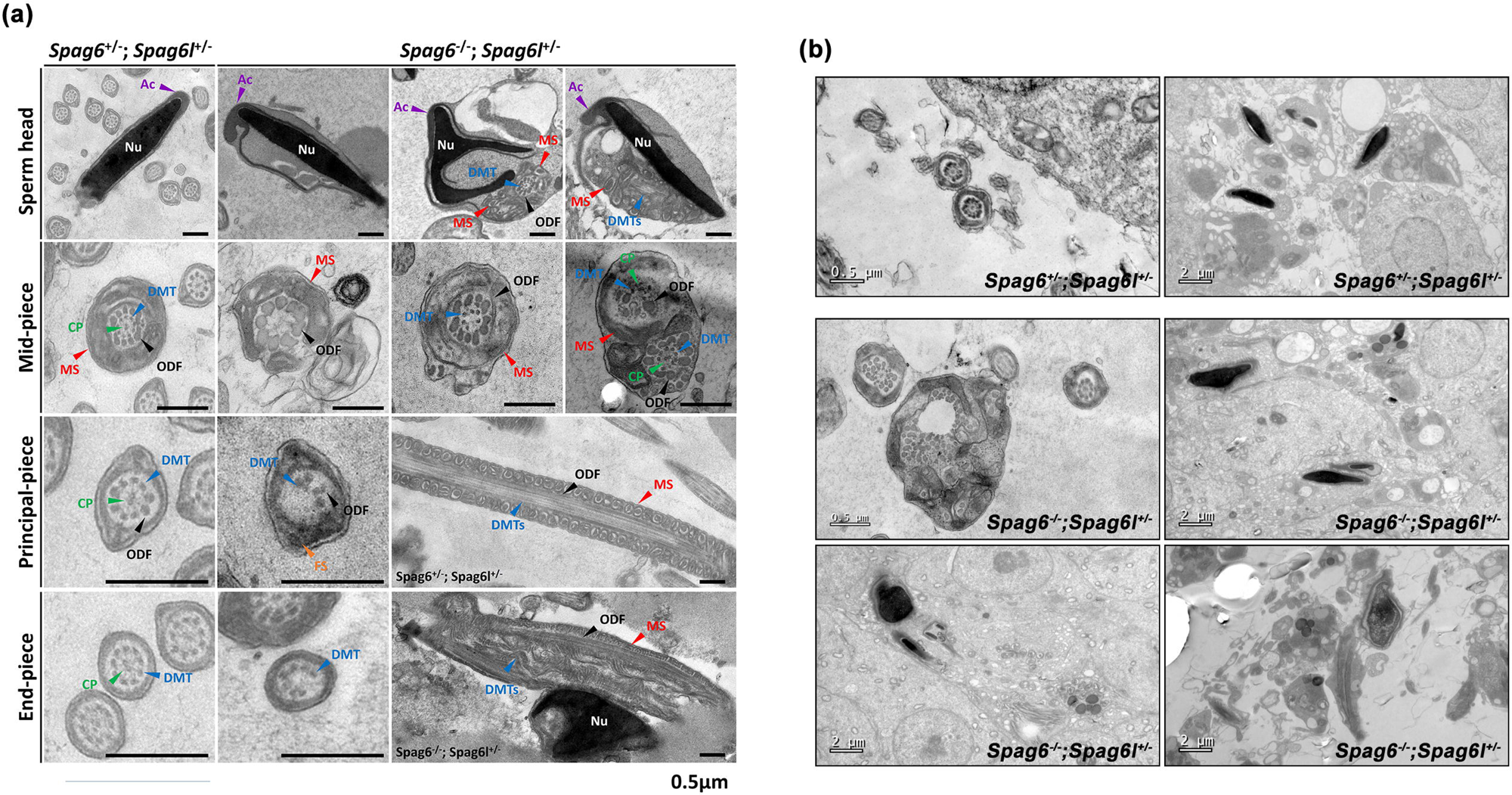
Ultrastructural changes in epididymal sperm and spermatogenic cells from *Spag6^+/−^_;_ Spag6l^+/−^* and *Spag6^−/−^_;_ Spag6l^+/−^* mice. **(a)** Epididymal sperm. Normal chromatin structure, ‘9+2’ axonemal microtubule cores and peri-axonemal structures were present in *Spag6^+/−^_;_ Spag6l^+/−^* mice. Malformed sperm heads with abnormal acrosome and residual cytoplasmic remnants were observed in *Spag6^−/−^_;_ Spag6l^+/−^* mice. Partial or complete loss of the ‘9+2’ microtubule arrangement, disorganized outer dense fibers, distorted mitochondrial sheath and fibrous sheath were also observed in sperm flagella from *Spag6^−/−^_;_ Spag6l^+/−^* mice. **(b)** Seminiferous tubules. Spermatids from *Spag6^+/−^_;_ Spag6l^+/−^* mice displayed a well-organized axoneme and properly shaped sperm heads. Misshapen sperm heads and disorganized axoneme and accessory structures of sperm flagella were observed in *Spag6^−/−^_;_ Spag6l^+/−^* spermatids.

The ultrastructure of the testicular spermatids was also examined. Similar to epididymal spermatozoa, TEM analysis revealed well-organized axonemes, intact accessory structures, and properly shaped sperm heads in the *Spag6^+/−^_;_ Spag6l^+/−^* spermatids. However, misshapen sperm heads and disorganized axoneme and accessory structures of sperm flagella were observed in the *Spag6^−/−^ Spag6l^+/−^* spermatids (**Figure 5b**), confirming that the ultrastructural abnormalities in the epididymal sperm originate during spermiogenesis.

### SPAG6 and SPAG6L exhibit differential binding affinity to SPINK2

Our previous study demonstrated that SPAG6L interacts with SPINK2, and that the expression of SPINK2 was significantly reduced in *Spag6l* knockout mice (8). To test the interaction between SPAG6 and SPINK2, a direct yeast two-hybrid experiment was performed, and we confirmed that SPAG6 also binds to SPINK2 (**Figure 6a**). Moreover, a protein complementation assay (PCA) was performed to compare the binding capacities of SPAG6 and SPAG6L for SPINK2. Constructs encoding SPAG6, SPAG6L, or SPINK2 fused to either N-terminal (N-Luc) or C-terminal (C-Luc) domains of luciferase were co-transfected into HEK293T cells, and luciferase activity was quantified. SPAG6 exhibited significantly stronger binding affinity to SPINK2 than SPAG6L, regardless of the fusion configuration (N-Luc/SPAG6 + SPINK2/C-Luc or N-Luc/SPINK2 + SPAG6/C-Luc). These results indicate that the evolutionarily divergent SPAG6L exhibits reduced binding capacity for SPINK2 (**Figure 6b**).

**Figure 6.**
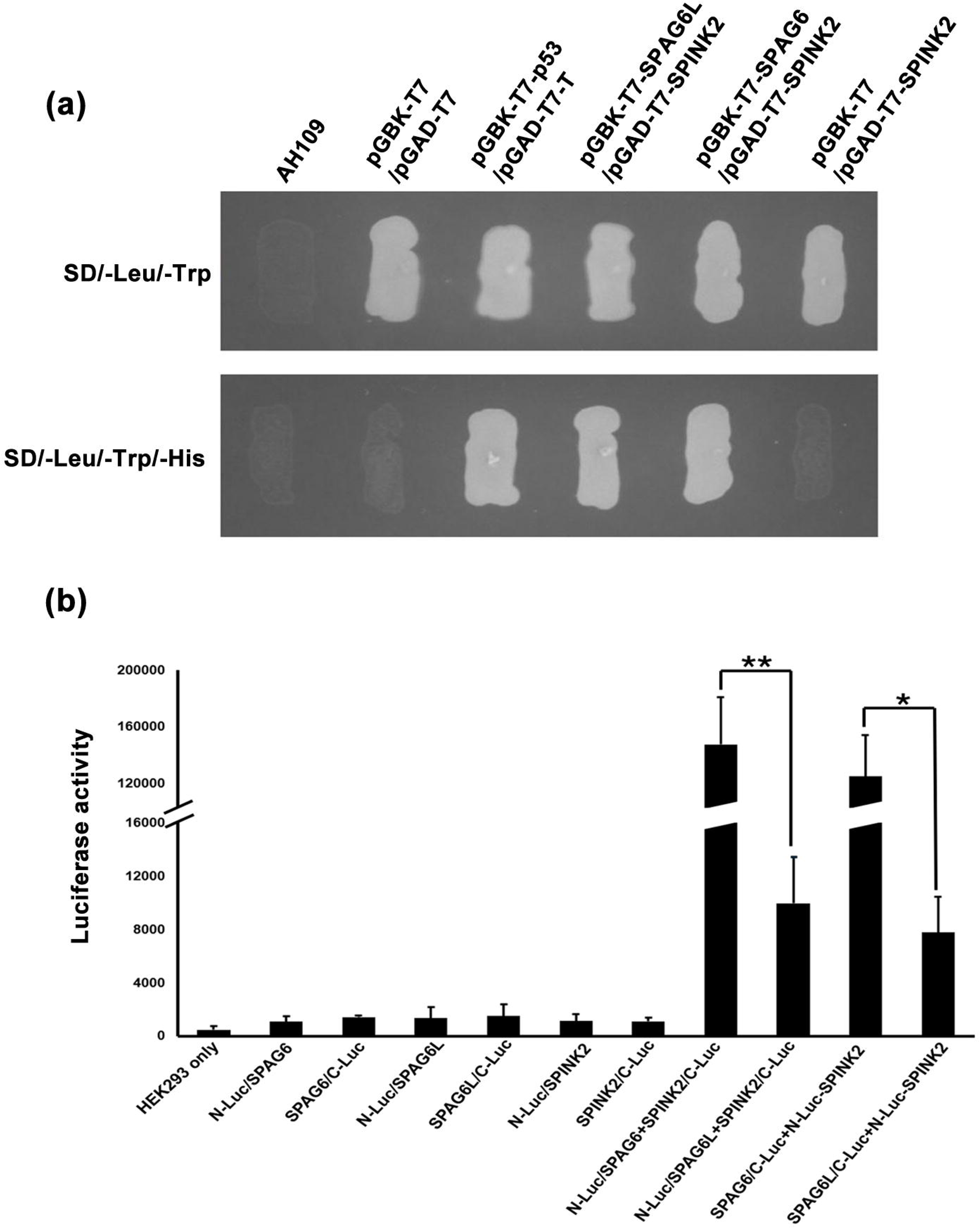
Differential binding affinity of SPINK2 with SPAG6 and SPAG6L. **(a)** SPINK2 interacts with SPAG6 or SPAG6L in yeast. Pairs of indicated plasmids were co-transformed into AH109 yeast, and the transformed yeast were grown on either selection plates (lacking tryptophan, leucine, and histidine) or non-selection plates (lacking tryptophan and leucine). Yeast expressing SPAG6/SPINK2, SPAG6L/SPINK2 and p53/large T antigen pairs grew on selection plate. **(b)** Binding capacity of SPAG6 and SPAG6L for SPINK2 was measured by protein complementation assays. SPAG6, SPAG6L, or SPINK2 was fused to either N-terminal (N-Luc) or C-terminal (C-Luc) domains of luciferase. The indicated plasmids were co-transfected into HEK293T cells, and luciferase activity were measured. SPAG6 exhibited significantly stronger binding to SPINK2 than SPAG6L.

SPINK2 is located in the acrosome of spermatids and spermatozoa. We performed immunofluorescence staining to analyze its localization at various stages of the spermatogenic cycle. In *Spag6^−/−^_;_ Spag6l^+/−^* mice, the SPINK2 signal was predominantly localized to the acrosome, exhibiting stage-specific expression patterns that coincided with acrosomal development. It was colocalized with PNA (a marker for the acrosome) and showed particularly intense staining at the apical tip of the acrosome. Despite the presence of abnormal sperm heads and malformed acrosomes in the *Spag6^−/−^_;_ Spag6l^+/−^* mice, the SPINK2 signal still overlapped with PNA staining, restricted to the acrosomal region (**Figure 7a**). Furthermore, Western blot analysis revealed a significant reduction in the testicular SPINK2 protein level in the *Spag6^−/−^_;_ Spag6l^+/−^* mice compared to the *Spag6^+/−^_;_ Spag6l^+/−^* mice (**Figure 7b**).

**Figure 7.**
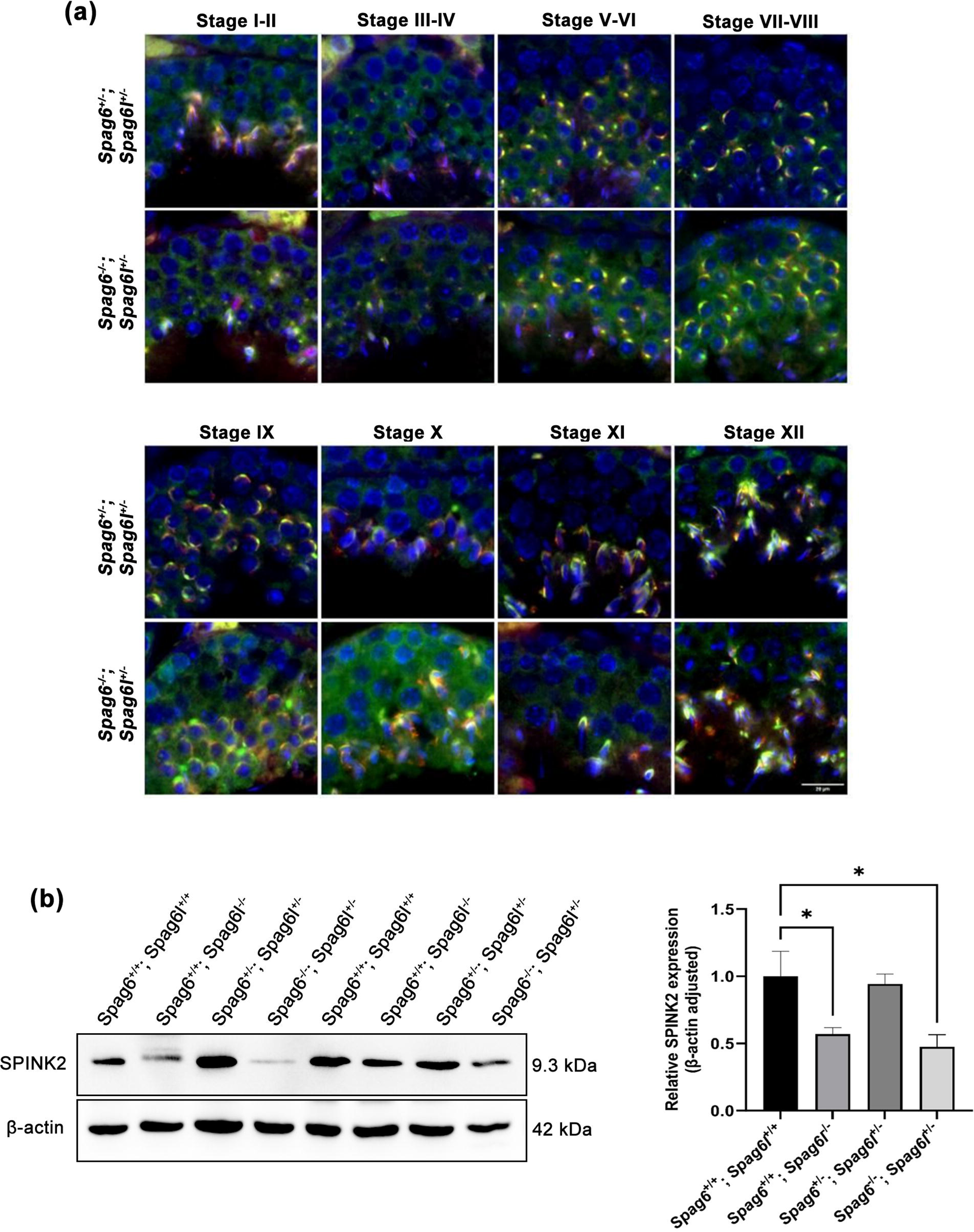
Localization and expression of SPINK2 in the seminiferous tubules of *Spag6^−/−^_;_ Spag6l^+/−^* mice. **(a)** The localization of SPINK2 in seminiferous tubules was analyzed by immunofluorescence microscopy. In Stages I-IV, SPINK2 was predominantly localized to the acrosomal region of round and elongating spermatids; in Stages V-VIII, while maintaining strong acrosomal expression, SPINK2 showed increased cytoplasmic accumulation in round spermatids; and throughout Stages IX-XII its distribution dynamically followed acrosomal maturation along the nuclear surface of elongated spermatids. Notably, SPINK2 signals consistently colocalized with PNA staining across all developmental stages (I-XII). In *Spag6^−/−^_;_ Spag6l^+/−^* mice, SPINK2 localization patterns remained comparable to *Spag6^+/−^_;_ Spag6l^+/−^* during Stages I-VIII, preserving both acrosomal predominance and PNA colocalization. However, during Stages IX-XII, as mutant spermatids developed abnormal head morphology, SPINK2 distribution became progressively distorted yet remarkably maintained its association with PNA-positive acrosomal structures despite these morphological aberrations. **(b)** The testicular expression of SPINK2. The expression levels of SPINK2 were analyzed by Western blot. The expression of SPINK2 in the *Spag6^+/+^_;_ Spag6l^−/−^* mice was lower than those in the *Spag6^+/+^_;_ Spag6l^+/+^* mice. Similarly, its expression was lower in the *Spag6^−/−^_;_ Spag6l^+/−^* mice, compared to the *Spag6^+/−^_;_ Spag6l^+/−^* mice.

### Abnormal levels of sperm flagellar proteins in *Spag6^−/−^_;_ Spag6l^+/−^* mouse testes

Given abnormal sperm flagellar ultrastructure in the *Spag6^−/−^_;_ Spag6l^+/−^* mice, we further examined testicular levels of AKAP4 and SPAG16, key components of the fibrous sheath and axoneme, respectively. APAP4 is initially translated as a 110 kDa pro-AKAP4 and the processed 84 kDa protein is incorporated into fibrous sheath during sperm flagella formation (21). In both surviving *Spag6^+/+^_;_ Spag6l*^−/−^ and *Spag6^−/−^_;_ Spag6l^+/−^* mice, the AKAP4/pro-AKAP4 ratio was significantly reduced compared to the *Spag6^+/−^_;_ Spag6l^+/−^* mice (**Figure 8a**). We previously reported that the full length testicular SPAG16 level was reduced in the *Spag6l*^−/−^ mice (22). In the *Spag6^−/−^_;_ Spag6l^+/−^* mice, testicular SPAG16 is also significantly reduced (**Figure 8b**).

**Figure 8.**
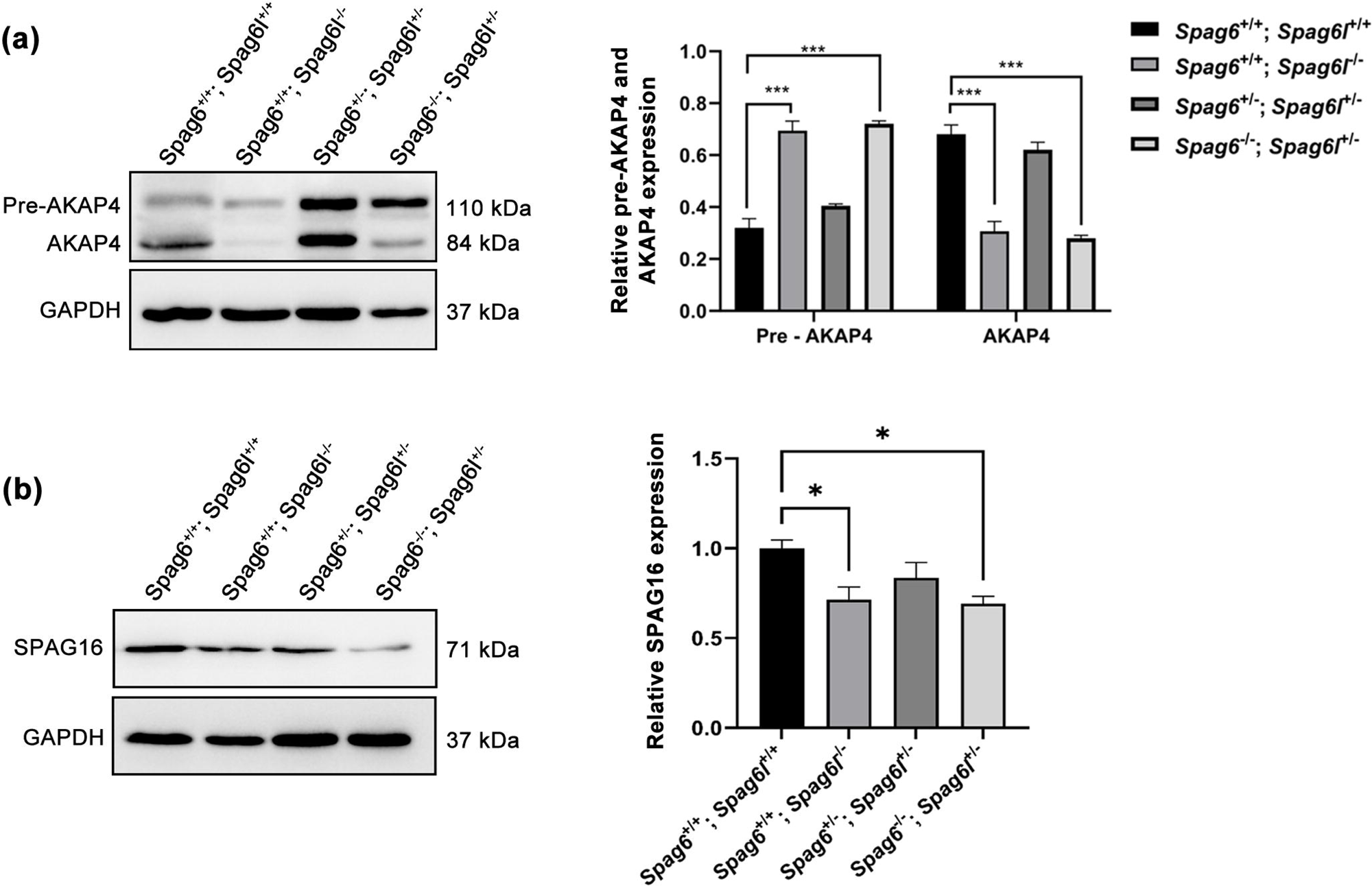
Abnormal testicular AKAP4 and SPAG16 levels in the *Spag6^−/−^_;_ Spag6l^+/−^* mice. **(a)** Examination of testicular AKAP4 levels. Left: a representative Western blot result; right: quantitative analysis of AKAP4. In the *Spag6^+/+^_;_ Spag6l^−/−^* and *Spag6^−/−^_;_ Spag6l^+/−^* mice, the processed 84 kDa AKAP4 levels were significantly reduced. Data are expressed as means ± SD (n = 4). \*\*\**P* < 0.001 compared with the *Spag6^+/+^_;_ Spag6l^+/+^* mice. **(b)** Examination of testicular SPAG16 levels. Left: a representative Western blot result; right: quantitative analysis of SPAG16. Like in the *Spag6^+/+^_;_ Spag6l^−/−^* mice, the 71 kDa SPAG16 level was significantly reduced in the *Spag6^−/−^_;_ Spag6l^+/−^* mice. Data are expressed as means ± SD (n = 4). \**P* < 0.05 compared with the *Spag6^+/+^_;_ Spag6l^+/+^* mice.

### Abnormal tubulin acetylation in *Spag6^−/−^_;_ Spag6l^+/−^* mouse testes

Post-translational modifications of tubulin play essential roles in regulating microtubule functions in cilia and flagella (23). Our previous study demonstrated that overexpression of *Spag6l* increased the expression of acetylated tubulin, whereas *Spag6l* deficiency markedly decreased it in mouse embryonic fibroblasts (13). We therefore assessed the levels of acetylated α-tubulin in *Spag6* and *Spag6l* knockout mice. Consistent with the previous cellular study, testicular levels of acetylated α-tubulin were significantly reduced in the *Spag6l^−/−^* mice (**Figure 9a**). Intriguingly, acetylated α-tubulin were markedly elevated in the *Spag6^−/−^* mice (**Figure 9b**). Further analysis of *Spag6^−/−^ Spag6l^+/−^* mice revealed intermediate acetylated α-tubulin levels between those of *Spag6^−/−^* and *Spag6l^−/−^* mice (**Figure 9c**). These findings suggest that SPAG6 and SPAG6L cooperatively regulate tubulin acetylation in testes

**Figure 9.**
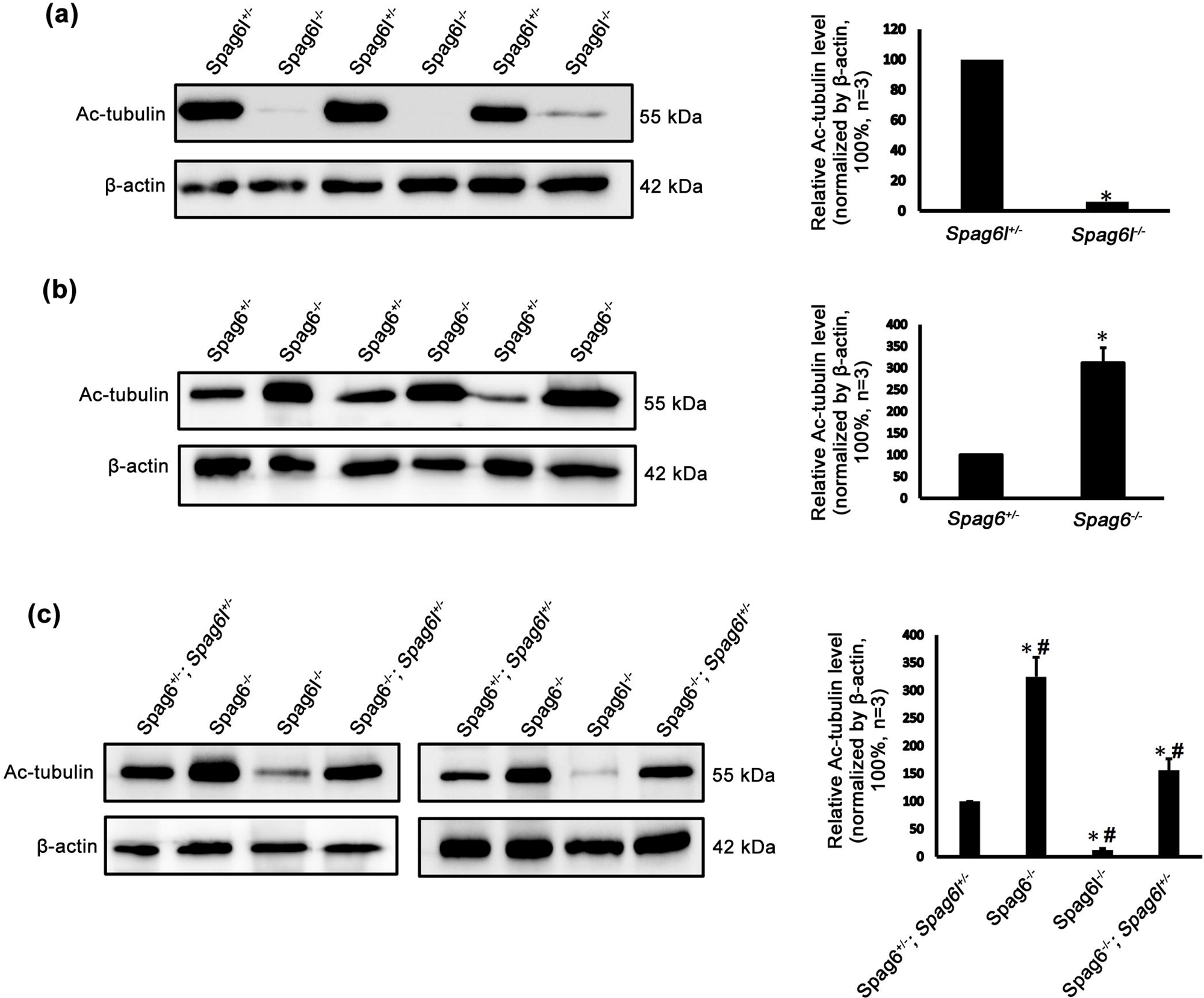
Abnormal tubulin acetylation in *Spag6^−/−^; Spag6l^+/−^* mice. **(a)** The level of acetylated α-tubulin in testes of *Spag6l^−/−^*mice was significantly downregulated, compared to the *Spag6l^+/−^* mice. **(b)** Acetylated α-tubulin level was markedly elevated in testes of *Spag6^−/−^* mice, compared to the *Spag6^+/−^* mice. **(c)** The level of acetylated α-tubulin in the testes of *Spag6^−/−^; Spag6l^+/−^* mice showed an intermediate expression level between that observed in *Spag6^−/−^* and *Spag6l^−/−^* mice.

## Discussion

Gene duplication is the primary source of new genes and is one of the key factors in driving speciation and phenotypic diversity. Although most duplicated genes undergo pseudogenization and are eventually eliminated, the retained duplicated genes often evolve asymmetrically, with one copy acquiring novel functions (24, 25). In mice, the *Spag6l* gene is believed to have evolved from the *Spag6* gene through a gene duplication event (6). *Spag6l^−/−^* mice exhibit multiple defects including hydrocephalus, hearing loss and male infertility, while *Spag6^−/−^* mice appear to be grossly normal (2, 7, 10, 26, 27). These findings suggest that *Spag6l* may have acquired additional functions during evolution, and that it plays a more prominent role in physiological processes than the ancestral *Spag6* gene.

In the present study, we generated compound *Spag6^+/−^_;_ Spag6l^+/−^* and *Spag6^−/−^_;_ Spag6l^+/−^* mice and explored the reproductive phenotypes in males. While the fertility and sperm parameters of *Spag6^+/−^_;_ Spag6l^+/−^* mice were grossly normal, *Spag6^−/−^_;_ Spag6l^+/−^* mutant males were completely infertile. These findings suggest that a single functional *Spag6l* allele is sufficient for normal spermatogenesis when at least one *Spag6* allele is present, and that there is partial functional compensation of SPAG6 for SPAG6L haploinsufficiency. Although all germ cell types were observed in the seminiferous tubules of *Spag6^−/−^_;_ Spag6l^+/−^* mice, morphological abnormalities were detected in the acrosome, manchette and flagella. Thus, male infertility of *Spag6^−/−^_;_ Spag6l^+/−^*mice may result from defective spermiogenesis.

Our previous study showed that *Spag6^−/−^* sperm exhibit normal morphology and function (7). In contrast, *Spag6^−/−^_;_ Spag6l^+/−^* sperm display severe disorganization of axoneme and peri-axonemal structures and reduced sperm motility, which are likewise observed in *Spag6l^−/−^* sperm (2). This suggests that SPAG6, which likely shares functional redundancy with SPAG6L, also mediates flagella assembly only when sufficient gene product of *Spag6l* is maintained. These structural and functional abnormalities of *Spag6^−/−^_;_ Spag6l^+/−^* sperm may result from impaired intraflagellar transport (IFT), as previously reported in flagella of IFT-deficient mice (28–30). In our yeast two-hybrid screen using SPAG6L as bait, IFT140 was identified as a binding partner (8), indicating a role for SPAG6L in IFT. Although the interaction between SPAG6 and IFT proteins remained uncharacterized, the high similarity in amino acid sequences between SPAG6 and SPAG6L suggests functional conservation, indicating SPAG6 is likely involved in the IFT machinery.

Previous studies demonstrated that the acrosomal protein, SPINK2, inhibits the protease activity of acrosin, and knockout of *Spink2* disrupts acrosome biogenesis in mice (31, 32). We previously demonstrated that SPAG6L interacts with SPINK2 and is likely to stabilize SPINK2 *in vivo* (8). This study further revealed that SPAG6 also binds SPINK2 with higher affinity than SPAG6L. While normal acrosome morphology was observed in both *Spag6^−/−^* and *Spag6l^−/−^* mice (7, 8), testicular SPINK2 levels were dramatically decreased and acrosomes were malformed in *Spag6^−/−^_;_ Spag6l^+/−^* mice. These findings suggest that SPAG6 and SPAG6L may functionally compensate for each other and cooperatively regulate SPINK2 activity and acrosome biogenesis. The significantly reduced testicular mature AKAP4 and SPAG16 levels might be another factor contributing to the abnormal sperm flagellar structure.

The manchette is a transient microtubule-based structure critical for spermiogenesis. It orchestrates sperm nuclear shaping and mediates cargo transport essential for flagellar development (33, 34). Microtubules, evolutionarily conserved heterodimers composed of α- and β-tubulin, play critical roles in eukaryotic cells (35). Post-translational modifications (PTMs) of tubulins are thought to regulate microtubule functions in specialized organelles (36). Notably, manchette and axoneme microtubules demonstrate characteristic PTMs including acetylation, glutamylation, and tyrosination, which dynamically regulate microtubule stability and interaction of microtubule-associated proteins during spermiogenesis (37, 38). This study revealed significant dysregulation of acetylated α-tubulin levels across different knockout models. Testicular acetylated α-tubulin displayed opposite expression between *Spag6l^−/−^* and *Spag6^−/−^* mice, while the levels of acetylated α-tubulin showed intermediate expression profiles in *Spag6^−/−^_;_ Spag6l^+/−^* mice. The results suggest potential antagonistic functions between SPAG6 and SPAG6L in modulating tubulin acetylation during spermiogenesis. The dysregulated tubulin acetylation in *Spag6^−/−^_;_ Spag6l^+/−^* spermatids likely contributes to abnormal formation of the manchette and sperm flagella. In mammals, microtubule acetylation is regulated by α-tubulin acetyltransferase 1 (α-TAT1) and histone deacetylase 6 (HDAC6), which catalyze tubulin acetylation and deacetylation, respectively (39). Knockout of α-TAT1 in mice abolishes detectable acetylation in flagella and impairs sperm motility (40). Similarly, decreased levels of acetylated α-tubulin were observed in sperm from human asthenozoospermia patients (41). Conversely, inhibition of HDAC6 activity in rat sperm compromises axonemal microtubule polymerization and reduces sperm motility (42). Given that both SPAG6 and SPAG6L contain eight contiguous armadillo domains that mediate protein-protein interaction (7), they may differentially regulate α-TAT1 and HDAC6 activity through protein interaction. However, whether SPAG6 and SPAG6L regulate spermiogenesis by modulating the expression, subcellular localization or activity of these enzymes remains to be elucidated.

## Acknowledgements

This research was supported by Wayne State University Start-up fund, NIH grants HD105944 (ZZ), HD107579 (ZZ), and HD114311 (ZZ). Aminata Touré is funded by INSERM, CNRS, Université Grenoble Alpes and the Agence Nationale pour la Recherche (ANR-17-CE13-0023 DIVERCIL, ANR-19-CE17-0014 FLAGELOME).

## Data Availability Statement

The data is available from the corresponding author upon reasonable request.

## Author Contributions

Y. Liu, W. Li, T. Li, C. Zheng, C. Niu, A. Schmitt, Y. Yap, M. Abdulghani performed research; S. Yuan, JF Strauss III, A Toure contributed to data analyses; C Melander, Y. Liu, L. Zhang, A Toure and Z Zhang wrote and proofread the paper.

## Supplementary Information

**Figure S1. Normal development of *Spag6^−/−^; Spag6l^+/−^* mice.**

Representative gross images of 3-month-old *Spag6^−/−^; Spag6l^+/−^* mice and littermates. No difference was observed between these mice.

**Figure S2. Morphological examination of epididymal sperm by light microscopy at low magnification.** Sperm density of the *Spag6^+/−^_;_ Spag6l^+/−^* mice is significantly higher than observed in the *Spag6^−/−^_;_ Spag6l^+/−^* mice at the same dilution.

**Figure S3. More ultrastructural images of principle piece of the epididymal sperm in the *Spag6^+/−^_;_ Spag6l^+/−^* and *Spag6^−/−^; Spag6l^+/−^* mice.**

Notice that the two longitudinal columns are associated with microtubule doublets 3 and 8, and the two semicircumferential ribs are symmetrical in the *Spag6^+/−^_;_ Spag6l^+/−^* mice. However, in the *Spag6^−/−^; Spag6l^+/−^* mice, the two semicircumferential ribs showing defective or asymmetric organization, and some sperm had three ribs.

## References

1. Smith EF, Lefebvre PA. The role of central apparatus components in flagellar motility and microtubule assembly. Cell Motil Cytoskeleton. 1997. 38(1): 1–8.

2. Sapiro R, Kostetskii I, Olds-Clarke P, Gerton GL, Radice GL, Strauss III JF. Male infertility, impaired sperm motility, and hydrocephalus in mice deficient in sperm-associated antigen 6. Mol Cell Biol. 2002. 22(17): 6298–305.

3. Neilson LI, Schneider PA, Van Deerlin PG, Kiriakidou M, Driscoll DA, Pellegrini MC, et al. cDNA cloning and characterization of a human sperm antigen (SPAG6) with homology to the product of the chlamydomonas PF16 locus. Genomics. 1999. 60(3): 272–80.

4. Khan MR, Akbari A, Nicholas TJ, Castillo-Madeen H, Ajmal M, Haq TU, et al. Genome sequencing of pakistani families with Male infertility identifies deleterious genotypes in SPAG6, CCDC9, TKTL1, TUBA3C, and M1AP. Andrology. 2025 13(5):1093–1104.

5. Sapiro R, Tarantino LM, Velazquez F, Kiriakidou M, Hecht NB, Bucan M, et al. Sperm antigen 6 is the murine homologue of the Chlamydomonas reinhardtii central apparatus protein encoded by the PF16 locus. Biol Reprod. 2000. 62(3): 511–8.

6. Qiu H, Gołas A, Grzmil P, Wojnowski L. Lineage-specific duplications of muroidea faim and Spag6 genes and atypical accelerated evolution of the parental Spag6 gene. J Mol Evol. 2013. 7(3):119–29.

7. Yap YT, Li W, Zhou Q, Haj-Diab S, Chowdhury DD, Vaishnav A, et al. The ancient and evolved mouse sperm-associated antigen 6 genes have different biologic functions In vivo. Cells. 2022. 11(3):336.

8. Liu Y, Zhang L, Li W, Huang Q, Yuan S, Li Y, et al. The sperm-associated antigen 6 interactome and its role in spermatogenesis. Reprod (camb Engl). 2019. 158(2):181–97.

9. Alciaturi J, Anesetti G, Irigoin F, Skowronek F, Sapiro R. Distribution of sperm antigen 6 (SPAG6) and 16 (SPAG16) in mouse ciliated and non-ciliated tissues. J Mol Histol. 2019. 50(3):189–202.

10. Li X, Zhang D, Xu L, Han Y, Liu W, Li W, et al. Planar cell polarity defects and hearing loss in sperm-associated antigen 6 (Spag6)-deficient mice. Am J Physiol, Cell Physiol. 2021. 320(1): C132–41.

11. Cooley LF, El Shikh ME, Li W, Keim RC, Zhang Z, Strauss JF, et al. Impaired immunological synapse in sperm associated antigen 6 (SPAG6) deficient mice. Sci Rep. 2016. 6: 25840.

12. Teves ME, Sears PR, Li W, Zhang Z, Tang W, van Reesema L, et al. Sperm-associated antigen 6 (SPAG6) deficiency and defects in ciliogenesis and cilia function: polarity, density, and beat. PLOS One. 2014. 9(10): e107271.

13. Li W, Mukherjee A, Wu J, Zhang L, Teves ME, Li H, et al. Sperm associated antigen 6 (SPAG6) regulates fibroblast cell growth, morphology, migration and ciliogenesis. Sci Rep. 2015. 5:16506.

14. Yan R, Hu X, Zhang Q, Song L, Zhang M, Zhang Y, et al. Spag6 negatively regulates neuronal migration during mouse brain development. J Mol Neurosci: MN. 2015. 57(4): 463–9.

15. Shanley L, Lear M, Davidson S, Ross R, MacKenzie A. Evidence for regulatory diversity and auto-regulation at the TAC1 locus in sensory neurones. J Neuroinflammation. 2011. 8:10.

16. Huang Q, Liu H, Zeng J, Li W, Zhang S, Zhang L, et al. COP9 signalosome complex subunit 5, an IFT20 binding partner, is essential to maintain male germ cell survival and acrosome biogenesis†. Biol Reprod. 2020. 102(1):233–47.

17. Zhang S, Liu Y, Huang Q, Yuan S, Liu H, Shi L, et al. Murine germ cell-specific disruption of Ift172 causes defects in spermiogenesis and Male fertility. Reprod (camb Engl). 2020. 159(4):409–21.

18. Cavarocchi E, Sayou C, Lorès P, Cazin C, Stouvenel L, El Khouri E, et al. Identification of IQCH as a calmodulin-associated protein required for sperm motility in humans. Iscience. 2023. 26(8):107354.

19. Huang Q, Li W, Zhou Q, Awasthi P, Cazin C, Yap Y, et al. Leucine zipper transcription factor-like 1 (LZTFL1), an intraflagellar transporter protein 27 (IFT27) associated protein, is required for normal sperm function and Male fertility. Dev Biol. 2021. 477:164–76.

20. Yap YT, Li W, Huang Q, Zhou Q, Zhang D, Sheng Y, et al. DNALI1 interacts with the MEIG1/PACRG complex within the manchette and is required for proper sperm flagellum assembly in mice. Elife. 2023. 12:e79620.

21. Johnson LR, Foster JA, Haig-Ladewig L, VanScoy H, Rubin CS, Moss SB, Gerton GL. Assembly of AKAP82, a protein kinase A anchor protein, into the fibrous sheath of mouse sperm. Dev Biol. 1997. 192(2):340–50.

22. Zhang Z, Sapiro R, Kapfhamer D, Bucan M, Bray J, Chennathukuzhi V, McNamara P, Curtis A, Zhang M, Blanchette-Mackie EJ, Strauss JF 3rd. A sperm-associated WD repeat protein orthologous to Chlamydomonas PF20 associates with Spag6, the mammalian orthologue of Chlamydomonas PF16. Mol Cell Biol. 2002. 22(22):7993–8004.

23. Wloga D, Joachimiak E, Louka P, Gaertig J. Posttranslational Modifications of Tubulin and Cilia. Cold Spring Harb Perspect Biol. 2017. 9(6): a028159.

24. Pegueroles C, Laurie S, Albà MM. Accelerated evolution after gene duplication: a time-dependent process affecting just one copy. Mol Biol Evol. 2013. 30(8):1830–42.

25. He X, Zhang J. Rapid subfunctionalization accompanied by prolonged and substantial neofunctionalization in duplicate gene evolution. Genetics. 2005. 169(2):1157–64.

26. Li X, Xu L, Li J, Li B, Bai X, Strauss JF, et al. Otitis media in sperm-associated antigen 6 (Spag6)-deficient mice. PLOS One. 2014. 9(11): e112879.

27. Li X, Zhang D, Xu L, Liu W, Zhang N, Strauss JF, et al. Sperm-associated antigen 6 (Spag6) mutation leads to vestibular dysfunction in mice. J Pharmacol Sci. 2021. 147(4):325–30.

28. Zhang Z, Li W, Zhang Y, Zhang L, Teves ME, Liu H, et al. Intraflagellar transport protein IFT20 is essential for Male fertility and spermiogenesis in mice. Mol Biol Cell. 2016. 27(23):3705–16.

29. Zhang Y, Liu H, Li W, Zhang Z, Shang X, Zhang D, et al. Intraflagellar transporter protein (IFT27), an IFT25 binding partner, is essential for Male fertility and spermiogenesis in mice. Dev Biol. 2017. 432(1):125–39.

30. Shi L, Zhou T, Huang Q, Zhang S, Li W, Zhang L, et al. Intraflagellar transport protein 74 is essential for spermatogenesis and Male fertility in mice†. Biol Reprod. 2019. 101(1):188–99.

31. Lee B, Park I, Jin S, Choi H, Kwon JT, Kim J, et al. Impaired spermatogenesis and fertility in mice carrying a mutation in the Spink2 gene expressed predominantly in testes. J Biol Chem. 2011. 286(33): 29108–17.

32. Kherraf ZE, Christou-Kent M, Karaouzene T, Amiri-Yekta A, Martinez G, Vargas AS, et al. SPINK2 deficiency causes infertility by inducing sperm defects in heterozygotes and azoospermia in homozygotes. EMBO Mol Med. 2017. 9(8):1132–49.

33. Wei YL, Yang WX. The acroframosome-acroplaxome-manchette axis may function in sperm head shaping and Male fertility. Gene. 2018. 660:28–40.

34. Lehti MS, Sironen A. Formation and function of the manchette and flagellum during spermatogenesis. Reprod (camb Engl). 2016. 151(4): R43–54.

35. Goodson HV, Jonasson EM. Microtubules and microtubule-associated proteins. Cold Spring Harbor Perspect Biol. 2018. 10(6): a022608.

36. Konno A, Setou M, Ikegami K. Ciliary and flagellar structure and function--their regulations by posttranslational modifications of axonemal tubulin. Int Rev Cell Mol Biol. 2012. 294:133–70.

37. Kierszenbaum AL, Rivkin E, Tres LL. Cytoskeletal track selection during cargo transport in spermatids is relevant to Male fertility. Spermatogenesis. 2011. 1(3):221–30.

38. Mochida K, Tres LL, Kierszenbaum AL. Structural and biochemical features of fractionated spermatid manchettes and sperm axonemes of the azh/azh mutant mouse. Mol Reprod Dev. 1999. 52(4):434–44.

39. Donker L, Godinho SA. Rethinking tubulin acetylation: from regulation to cellular adaptation. Curr Opin Cell Biol. 2025. 94:102512.

40. Kalebic N, Sorrentino S, Perlas E, Bolasco G, Martinez C, Heppenstall PA. αTAT1 is the major α-tubulin acetyltransferase in mice. Nat Commun. 2013. 4:1962.

41. Bhagwat S, Dalvi V, Chandrasekhar D, Matthew T, Acharya K, Gajbhiye R, et al. Acetylated α-tubulin is reduced in individuals with poor sperm motility. Fertil Steril. 2014. 101(1):95–104.e3.

42. Chawan V, Yevate S, Gajbhiye R, Kulkarni V, Parte P. Acetylation/deacetylation and microtubule associated proteins influence flagellar axonemal stability and sperm motility. Biosci Rep. 2020. 40(12):BSR20202442.

