## Supplemental Table for "Two *Spag6* genes control sperm formation and male fertility in mice"

**Supplemental Table 1. The sequences of primers used for cloning.**

| **Primer name** | **Primer sequences (5’→3’)** | **Usage** |
| --- | --- | --- |
| Spag6-C-Luc-F | AAGCTTCGATGAGCCAGCGGCAGGTGCTGCAAG (Hind III) | N-Luc/*Spag6* *Spag6/*C-Luc |
| Spag6-N-Luc-F | AAGCTTATGAGCCAGCGGCAGGTGCTGCAAG (Hind III) |  |
| Spag6-Luc-R | ACCGGTGGGTTAATAAGAGGCTGATAGCTGTCG (Age I) |  |
| Spag6l-C-Luc-F | AAGCTTCGATGAGCCAGCGGCAGGTGCTGCAAG (Hind III) | N-Luc/*Spag6l*, *Spag6l/*C-Luc |
| Spag6l-N-Luc-F | AAGCTTATGAGCCAGCGGCAGGTGCTGCAAG (Hind III) |  |
| Spag6l-Luc-R | ACCGGTGGCAGTGGTTGGTAGCTGTCCACCCTC (Age I) |  |
| Spink2-N-Luc-F | AAGCTTATGCTGAGACTGGTGCTGTTGCT  (Hind III) | N-Luc/*Spink2*, *Spink2*/C-Luc |
| Spink2-C-Luc-F | AAGCTTCGATGCTGAGACTGGTGCTGTTGCT (Hind III) |  |
| Spink2-Luc-R | ACCGGTGGGCATGGCTCGTCTTTGATGATAT  (Age I) |  |

**Supplemental Table 2 Genotypes and phenotype of combined global *Spag6* & *Spag6l* mutant mice.**

| Genotype | *Spag6^+/+^; Spag6l^+/+^* | *Spag6^+/+^; Spag6l^+/-^* | *Spag6^+/+^; Spag6l^-/-^* | *Spag6^+/-^; Spag6l^+/+^* | *Spag6^+/-^; Spag6l^+/-^* | *Spag6^+/-^; Spag6l^-/-^* | *Spag6^-/-^; Spag6l^+/+^* | *Spag6^-/-^; Spag6l^+/-^* | *Spag6^-/-^; Spag6l^-/-^* |
| --- | --- | --- | --- | --- | --- | --- | --- | --- | --- |
| Phenotype | Normal | Normal | Infertile  and die | Normal | Normal | Infertile and die | Normal | Infertile | Die |
