## Supplementary figures and images for "Two *Spag6* genes control sperm formation and male fertility in mice"

### Figure S1

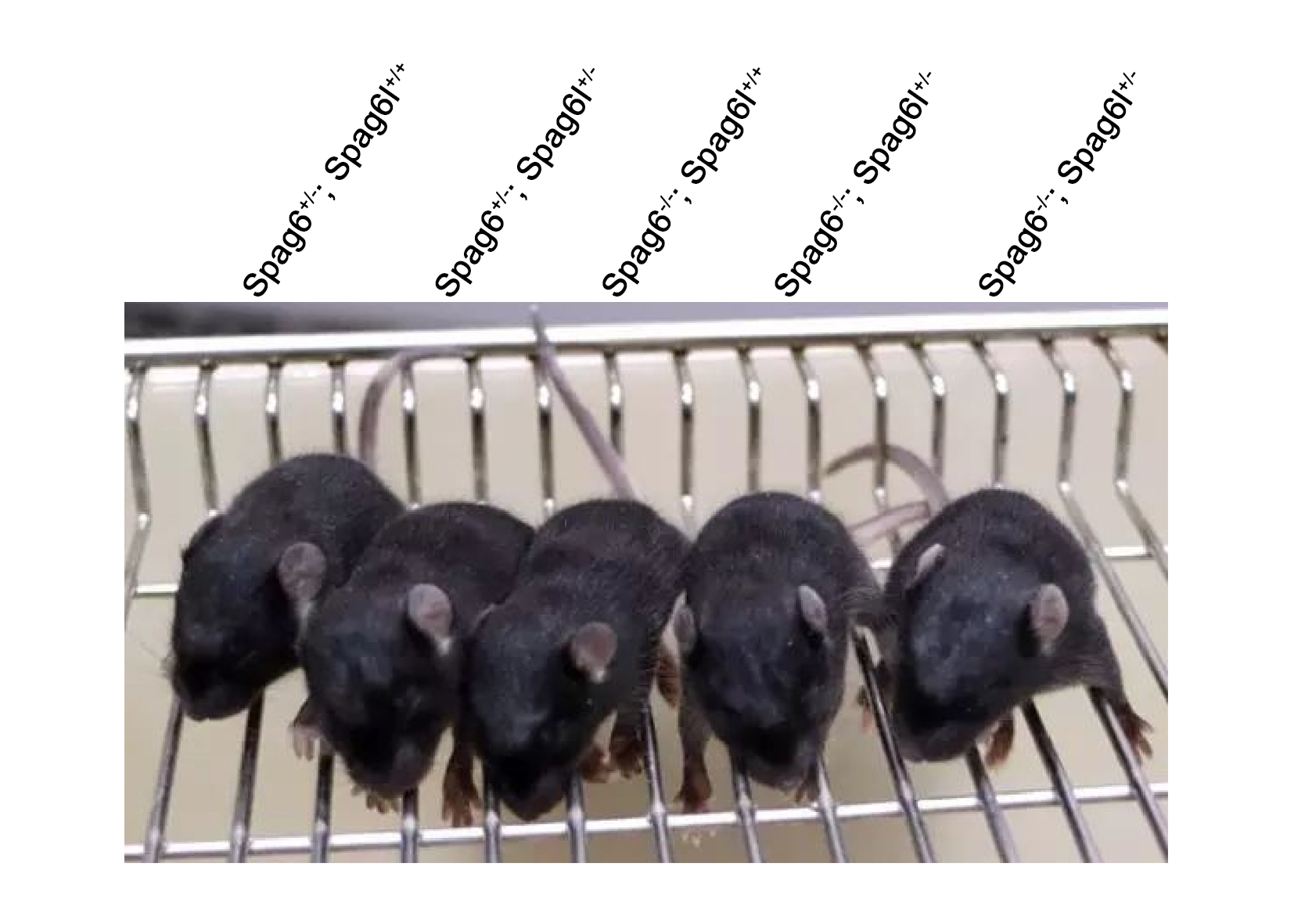

### Figure S2

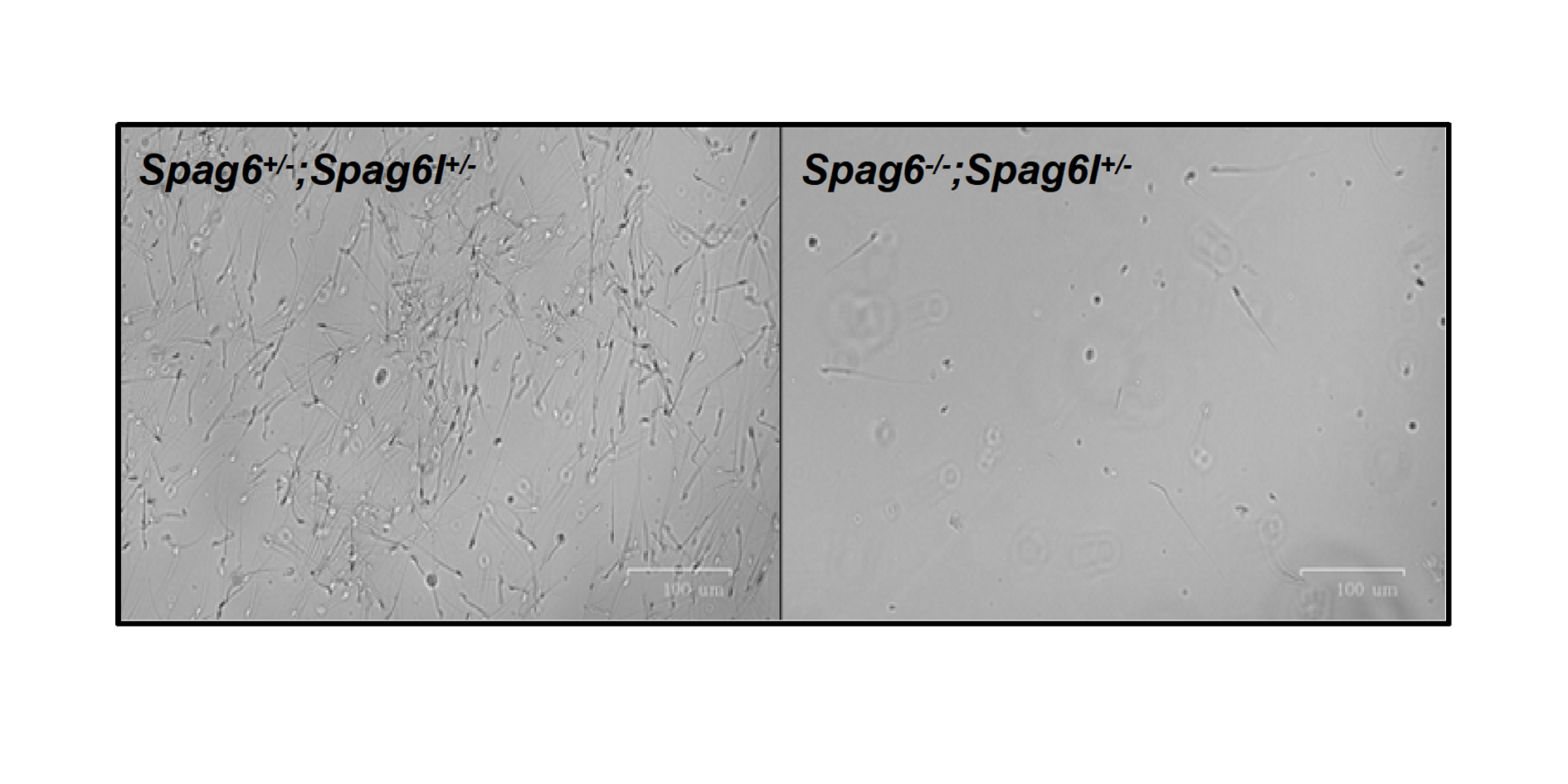

### Figure S3

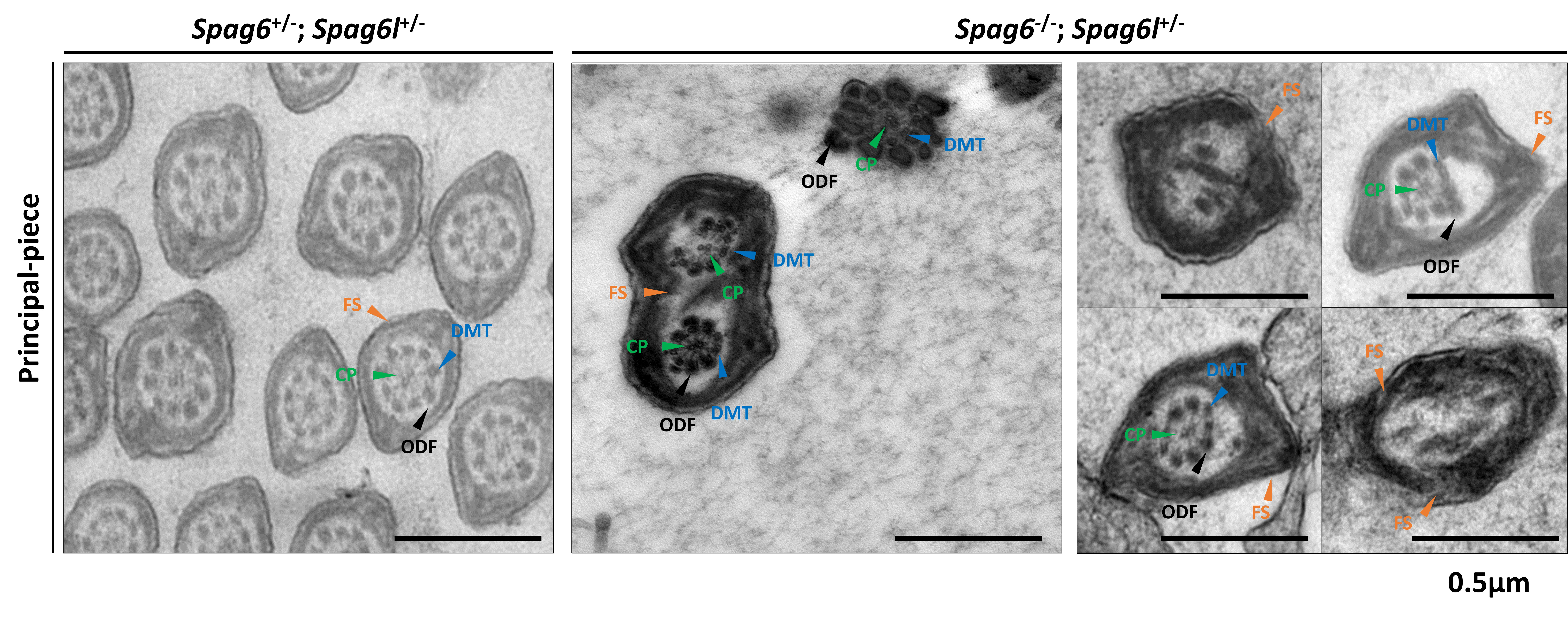
